# *Leptospira* transcriptome sequencing using long-read technology reveals unannotated transcripts and potential polyadenylation of mRNA molecules

**DOI:** 10.1101/2023.03.29.534852

**Authors:** Ruijie Xu, Dhani Prakoso, Liliana C. M. Salvador, Sreekumari Rajeev

**Affiliations:** Institute of Bioinformatics, University of Georgia, Athens, GA, 30602; Center for the Ecology of Infectious Diseases, University of Georgia, Athens, GA, 30602; Department of Biomedical and Diagnostic Sciences, College of Veterinary Medicine, University of Tennessee, Knoxville, TN, 37996, USA; Department of Infectious Diseases, College of Veterinary Medicine, University of Georgia, Athens, GA, 30602, USA

## Abstract

Leptospirosis caused by the spirochete bacteria *Leptospira* is an emerging zoonosis that causes life-threatening disease in humans and animals. However, a detailed understanding of *Leptospira’s* RNA profiles is limited. In this study, we sequenced and analyzed the transcriptome of multiple *Leptospira* serovars using Oxford Nanopore Technologies’ direct cDNA and direct RNA sequencing methods. We identified several operon transcriptional units, novel RNA coding regions, and evidence of potential posttranscriptional polyadenylation in the *Leptospira* transcriptome. Some coding regions of multiple RNA molecules that have not been previously annotated could be potential sRNA or ncRNA molecules that support gene expression regulation purposes in *Leptospira*. Many of the relative positions of these unannotated RNA coding regions were consistent in their neighboring coding regions across the reference genomes of two pathogenic *Leptospira* analyzed in this study. Besides, the majority of the unannotated coding regions and operon transcriptional units were not detected in the nonpathogenic *Leptospira*, suggesting potential virulence-related functions for these RNA molecules coding regions. Overall, our study confirms the utility of ONT’s sequencing in studying prokaryotic transcriptome profiles and offers a tool to improve our understanding of the structural composition of RNA molecules and prokaryotic polyadenylation. However, the findings from our study also warrant that the presence of homopolymers of adenine bases in the transcripts may interfere with the interpretation of bacterial transcriptome profiles. Carefully designed experiments are needed to unravel the role of the features described in this study in *Leptospira* virulence and pathogenesis.

**Author Summary:** Leptospirosis caused by the spirochete bacteria, *Leptospira,* is recognized as one of the most widespread zoonotic diseases. Leptospirosis is a neglected disease and has been estimated to cause over one million annual human clinical cases and over 60,000 deaths. *Leptospira* are maintained in the renal tubules of asymptomatic animal reservoirs, and in contaminated environments, such as soil and water. Studying gene expression profiles is an important component of bacterial pathogenesis studies, and a number of methods are available to do it. Our study compared various transcriptome sequencing methods using one of the latest next generation sequencing tools (Oxford Nanopore Technologies) and evaluated the compositions of RNA in different *Leptospira’s* transcriptomes. We identified many previously undescribed and potentially significant features within the *Leptospira* transcriptome that may contribute to the virulence and pathogenesis of *Leptospira sp*. Our findings lead to new opportunities for researchers to sequence prokaryotic RNA molecules and to unravel many regulatory mechanisms.

## INDRODUCTION

Leptospirosis caused by the spirochete bacteria, *Leptospira,* is recognized as one of the most widespread zoonotic diseases that can be equally fatal to humans and animals. Leptospirosis is a neglected disease and has been estimated to cause over one million annual human clinical cases and over 60,000 deaths [1]. Pathogenic *Leptospira* spp. are maintained in the renal tubules of asymptomatic animal reservoirs, and in contaminated environments, such as soil and water [2–4]. The members of the genus *Leptospira* are grouped into three distinct phylogenetic clades, pathogenic, intermediate, and saprophytic, based on their genetic similarities using DNA-DNA hybridization approach and 16S ribosomal RNA phylogeny [5,6]. Recent molecular classification extended *Leptospira* genus into two clades and four subclades (P1 and P2-pathogenic, and S1: and S2-saprophytic), using whole genome sequenced data [7]. Previous studies have found that the mechanisms responsible for *Leptospiral* pathogenesis and environmental/host adaptation were associated with the loss/acquisition of functional genes on the DNA level and changes in the overall gene expression profiles at the RNA level [8–12]. However, the current understanding of *Leptospiral* RNA regulation mechanisms is minimal.

The mechanisms of bacterial gene expression regulations that determine a wide range of physiological processes are now perceived to be more complex than previously thought [13]. The emergence of multiple RNA sequencing technologies has provided new opportunities to study gene expression patterns through the evaluation of bacterial RNA profiles [14]. Two long-read sequencing methods from PacBio’s single molecule real-time (SMRT) sequencing and Oxford Nanopore Technologies (ONT) sequencing have emerged as effective RNA sequencing tools that overcome some of the limitations of short-read sequencing [15,16]. While both technologies generate full-length RNA transcripts, ONT sequencing provides a more cost-efficient and portable option to directly sequence cDNA and RNA molecules without fragmentation and PCR amplification, alleviating sequencing biases associated with read-length and primer selection [17–19]. Moreover, the direct RNA sequencing (DRS) method from ONT has the ability to directly sequence the native RNA molecules without reverse transcribing RNA molecules into cDNA, which allows the preservation of posttranscriptional RNA modification markers [16,20] that play an important role in gene expression regulation mechanisms [21–23].

Prokaryote RNA profiles consist of a variety of molecules. Ribosomal RNAs (rRNA), which are responsible for the building of the protein-synthesizing organelle-ribosome, and transfer RNAs (tRNA), which are responsible for transporting amino acids to ribosomes for protein synthesis during translation [24–26], comprise over 95% of the total prokaryotic RNA profiles. On the other hand, the transcripts of messenger RNAs (mRNA), which contain the genetic codes for synthesizing functional proteins inside ribosomes during translation, only comprise less than 5% of the transcripts in the prokaryotic RNA profiles [24,25]. The ensemble of these RNA molecules regulates the pathogenesis and environmental adaption processes at the protein level [27,28]. In addition to the RNA molecules closely associated with the translation process above, the vast variety of noncoding RNA molecules (ncRNA) that does not go through translation are also an important component for regulating prokaryotic gene expressions by inhibit, activate, promote, or suppress the transcription/translation/degradation of RNA transcripts [29–31].

Since the posttranscriptional modification step of polyadenylation was not widely recognized in prokaryotes like in eukaryotic mRNA, to sequence prokaryotic mRNA, both ONT’s cDNA and DRS sequencing protocols require an enzymatic polyadenylation step to capture RNA transcripts during library preparation [32]. While experimenting with ONT’s RNA sequencing methods to evaluate *Leptospira* transcriptome, we accidentally skipped the polyadenylation step in our initial sequencing procedures and unexpectedly obtained abundant transcript reads. Recognizing that poly(A) tail in mRNA could be a feature in prokaryotic transcripts [22,23,33], we investigated the presence and composition of poly(A) stretches in *Leptospira* transcriptomes using ONT’s long-read sequencing methods. In this study, we conducted RNA-Seq experiments using two pathogenic and one nonpathogenic species with and without the polyadenylation step using ONT’s cDNA sequencing protocol. In addition, we performed DRS without polyadenylation on selected samples. Here, we reported the overall *Leptospira* RNA composition and characteristics utilizing ONT’s long-read sequencing method.

## RESULTS

### Overall sequencing and mapping results

A schematic representation of the sequencing process is shown in Fig S1. The total RNA from two pathogenic *L. interrogans* serovars, (Copenhageni [LIC] and Icterohemmorhagiae [LII]) and one nonpathogenic serovar of *L. biflexa* (serovar Patoc [LBP]) was sequenced with ONT’s cDNA protocol with and without additional enzymatic polyadenylation. Almost 3 million reads were sequenced from the six *Leptospira* cDNA samples used in this study with an average read length of ∼1,764 bp long. Over 95% of the sequenced cDNA reads of all samples had sequencing quality scores higher than 7 (QUAL >=7). The average read length was higher in samples (1,835 (SD:91)) with the polyadenylation (polyA) than in the samples (1,638 (SD: 115)) without the polyadenylation (nonpolyA). A summary statistic of cDNA sequences is shown in Table 1.

**Table 1.** Quality statistics of the *Leptospira raw* cDNA reads.

| Sample ID | Raw Read Statistics |  |  |  |  |  |  | Full-length (FL) Read Statistics |  |
| --- | --- | --- | --- | --- | --- | --- | --- | --- | --- |
|  | Number Reads | Read QUAL >= 7 | Cumulative Size(Gb) | Mean Read Len (bp) | N50 (bp) | Max Read Len (bp) | Number reads>1.kb | Number Reads | Mean Read Len (bp) |
| LIC-cDNA-nonpolyA | 199,615 | 184,279 | 0.32 | 1,601 | 2,230 | 24,092 | 139,679 | 29,821 | 520 |
| LIC-cDNA-polyA | 647,339 | 611,858 | 1.26 | 1,939 | 2,987 | 31,992 | 464,560 | 365,534 | 1,664 |
| LII-cDNA-nonpolyA | 317,151 | 292,752 | 0.49 | 1,547 | 2,167 | 17,330 | 214,681 | 41,000 | 433 |
| LII-cDNA-polyA | 512,875 | 480,989 | 0.92 | 1,799 | 2,841 | 47,547 | 351,270 | 249,622 | 1,514 |
| LBP-cDNA-nonpolyA | 516,304 | 503,805 | 1 | 1,928 | 2,877 | 20,570 | 420,667 | 40,006 | 746 |
| LBP-cDNA-polyA | 703,577 | 680,593 | 1.24 | 1,768 | 2,556 | 30,998 | 497,475 | 312,858 | 1,640 |

We extracted the full-length (FL) cDNA reads from each sample by determining the presence of forward (SSP) and reverse primers (VNP) within the individual raw reads (<u>epi2me-labs/pychopper: cDNA read preprocessing (github.com)</u>). Reads that failed to identify the presence of both primers were considered as read fragments and were filtered out. Then subsequences identified outside the SSP and VNP primers were trimmed from the raw reads along with the identified primers to obtain a FL reads dataset. After filtering, the FL cDNA reads consisted of, on average, around 10% of the reads in the raw non-polyadenylated samples and 50% of the reads in the raw polyadenylated samples (Table 1). After filtering and trimming, the average read length for the FL non-polyadenylated samples decreased by almost two-third compared to the raw non-polyadenylated samples (∼566 [SD: 162] bp long). In contrast, the average read length for the polyadenylated samples was only decreased by 12% (1,606 [SD: 81] bp long) (Table 1). The VNP primers hits in the non-polyadenylated reads presented lower average mapping quality to the VNP primers and with longer sequences trimmed off from the raw reads (trimmed-off lengths) during the FL read pre-processing step than the VNP primers hits’ mapping quality and trimmed-off lengths in the polyadenylated samples (Fig S2).

Over 99% of the raw cDNA reads were mapped to the corresponding reference genomes for both polyadenylated and non-polyadenylated samples. Whereas on average, 91% of the FL cDNA reads were mapped to the corresponding reference genomes (Fig 1a). The majority of the raw (94%) and FL (95%) cDNA reads in each sample were mapped to the coding regions of rRNAs. Only 5% and 3% of the raw and FL reads, respectively, mapped to the mRNA’s coding regions (Fig 1b). A higher mapping rate to mRNA coding regions was observed in both raw and FL cDNA reads from the polyadenylated samples compared to the non-polyadenylated ones. A higher number of distinct coding regions were mapped in the polyadenylated cDNA samples than in the non-polyadenylated ones, where FL cDNA reads aligned to half of the coding regions mapped by the raw cDNA reads in the polyadenylated samples and to less than one third in the non-polyadenylated ones (Fig 1c).

**Fig 1.**
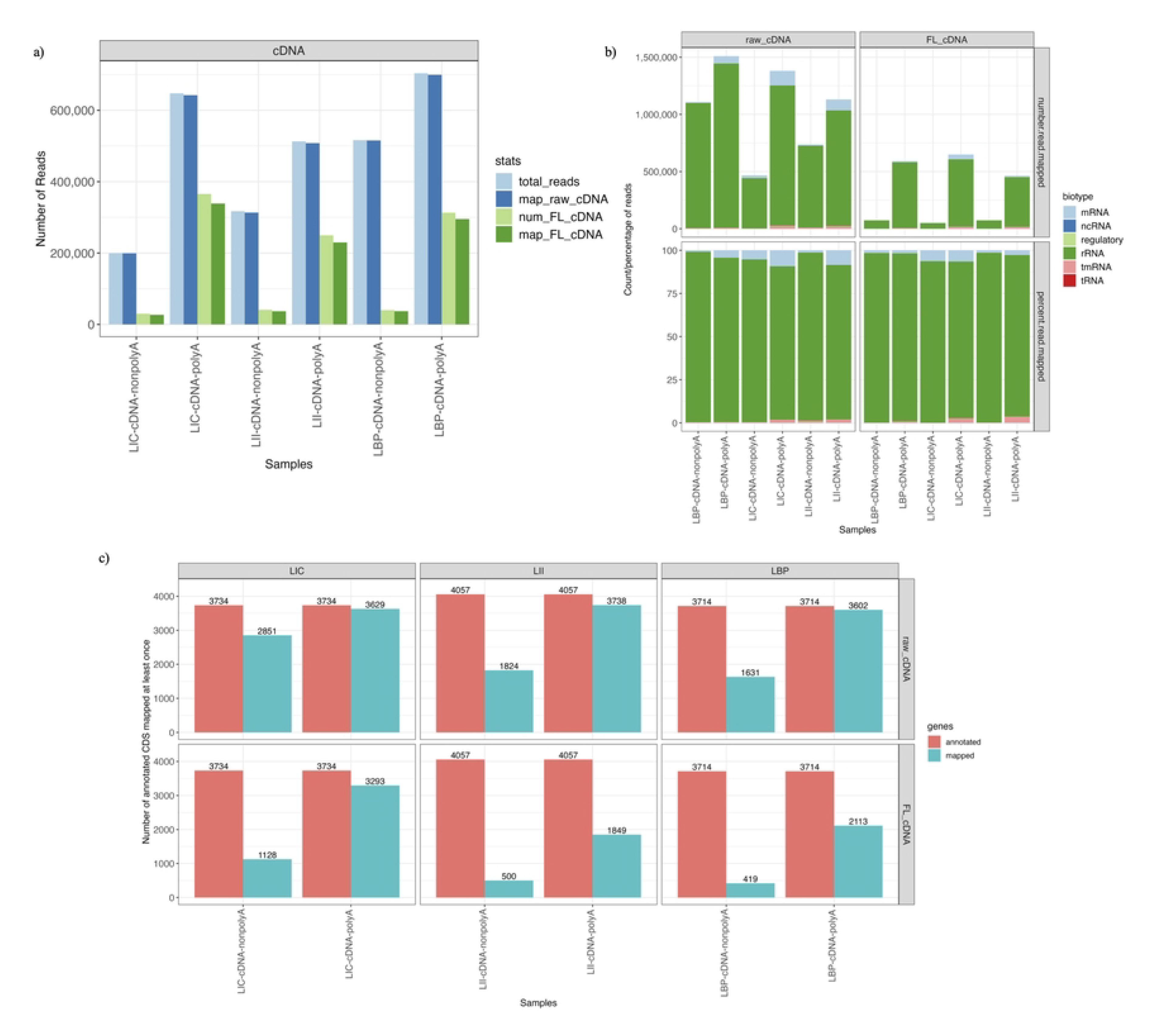
Mapping statistics of the cDNA reads. (a) The total number of raw (light blue) and FL (light green) cDNA reads sequenced within each sample and the number of raw (dark blue) and FL (dark green) reads mapped to their corresponding reference genomes. (b) The number (top) and percentage (bottom) of reads mapped to different RNA biotypes for raw (left) and FL cDNA reads (right). (c) The number of unique annotated coding regions mapped by raw cDNA (top row), FL cDNA (bottom row). Each column represents the samples obtained from a different serovar (LIC, LII, and LBP). The red bar on the left is the total number of annotated coding regions in the corresponding reference genomes and the blue bar on the right is the number of unique coding regions mapped at least once in the sample.

To further evaluate the presence and characteristics of the non-polyadenylated RNA fraction, we proceeded with ONT’s DRS sequencing protocol to sequence the original non-polyadenylated RNA molecules without reverse transcription on two technical replicates of the LIC serovar (LIC-DRS). The DRS samples showed a relatively high sequencing yield (Table 2) with an average read length of 767 and 868 bp, respectively, from each DRS sample. Over 90% of the sequenced reads had a quality score of over 7 (Table 2). Our attempt to sequence RNA by DRS after the polyadenylation step generated only less than ∼0.025 Gb of data, and these were not enough for a comparison analysis (data not shown).

**Table 2.**
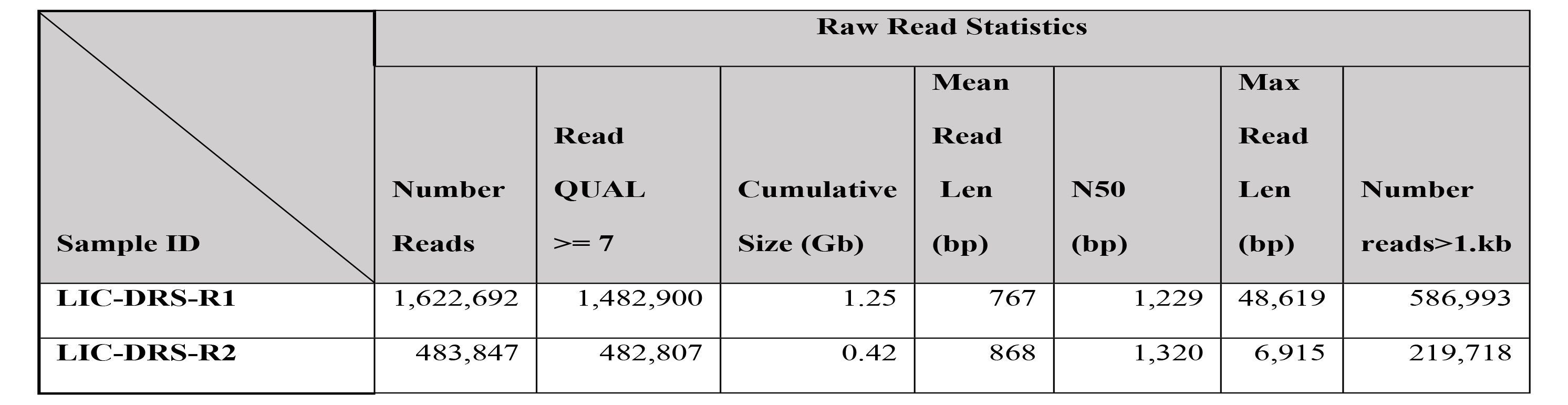
Raw DRS reads quality statistics.

| Sample ID | Raw Read Statistics |  |  |  |  |  |  |
| --- | --- | --- | --- | --- | --- | --- | --- |
| | Number Reads | Read QUAL $\geq 7$ | Cumulative Size (Gb) | Mean Read Len (bp) | N50 (bp) | Max Read Len (bp) | Number reads>1.kb |
| LIC-DRS-R1 | 1,622,692 | 1,482,900 | 1.25 | 767 | 1,229 | 48,619 | 586,993 |
| LIC-DRS-R2 | 483,847 | 482,807 | 0.42 | 868 | 1,320 | 6,915 | 219,718 |

An average of 36% of DRS reads mapped to the reference genome (Fig 2a) with an average mapping identity slightly lower (87%) than the cDNA reads (raw: 90%; FL: 93%). In contrast to the cDNA reads, over 71% of the DRS reads mapped to the coding regions of mRNA molecules, and only 28% of the reads mapped to the coding regions of rRNA molecules (Fig 2b). The DRS reads sequenced without polyadenylation aligned to more distinct annotated coding regions (Fig 2c) than the cDNA polyadenylated LIC sample (Fig 1c).

**Fig 2.**
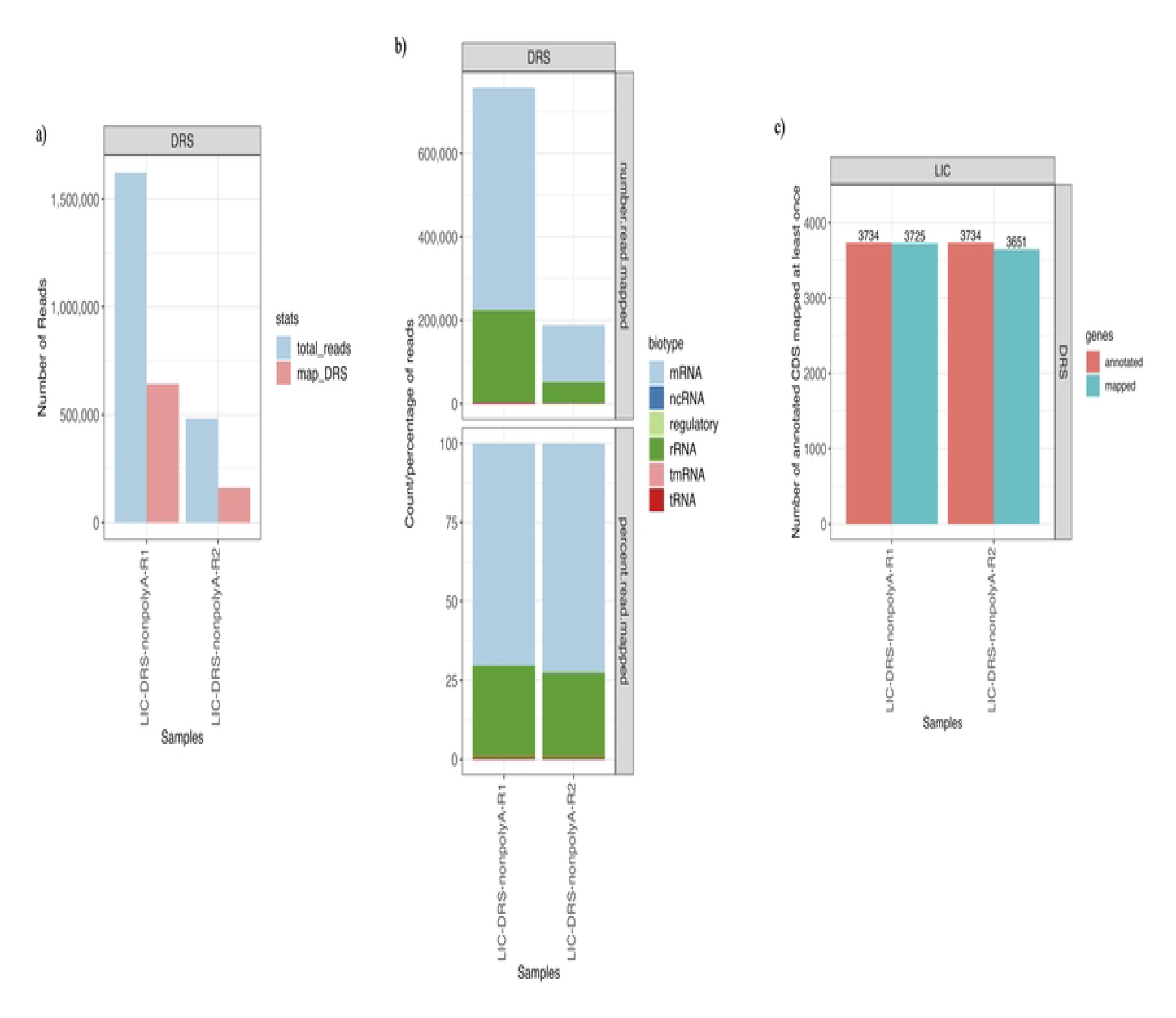
DRS samples’ reads and mapping statistics. (a) The number of DRS reads (light blue) sequenced within each sample and the number of DRS reads mapped to LIC’s reference genome (light red). (b) The number (top) and percentage (bottom) of reads mapped to different RNA biotypes within each DRS sample. (c) The number of distinct annotated coding regions mapped by DRS reads within each sample. The red bar on the left is the total number of annotated coding regions in the corresponding reference genomes and the blue bar on the right is the number of distinct coding regions mapped at least once in the sample.

### Relative mapping positions of individual reads

To assess the exact positions of the mapped reads relative to the coding regions of the RNA molecules, we classified the relative mapping positions of each read in relationship to its closest annotated coding region into six different categories (“complete”, “internal”, “noncoding”, “operons”, “partial (3’)” and “partial (5’)”) (Fig 3a). In general, only around 4% of the cDNA reads (both raw and FL) completely aligned to the annotated coding regions of their mapped RNA molecules (“complete”). From which, less than 0.5% of the non-polyadenylated cDNA reads (both raw and FL) were “complete” mapped reads. The largest “complete” mapped coding region in the non-polyadenylated samples was the 23S ribosomal RNA coding region (2,958 bp), and in the polyadenylated samples was the LIC12048 family lipoprotein coding region (4,509 bp). Most cDNA reads were found either embedding the annotated coding regions (“internal”) or covering only the 5’ (“partial (5’)”) or 3’ (“partial (3’)”) end of the coding regions of the mapped RNA molecules (Fig 3b), with more “partial (3’)” reads identified in the polyadenylated samples than in the non-polyadenylated ones in both raw and FL cDNA reads. Furthermore, we also observed that approximately 1% of the raw and FL cDNA reads mapped between two flanking RNA coding regions (“noncoding”), and that less than 1% of the raw and FL cDNA reads mapped across more than one RNA coding region (“operons”) (Fig 3b). To compare the RNA profiles of the cDNA reads with and without polyadenylation, we calculated the correlation between the coding regions coverage mapped for each sample between both fractions. The coding region coverage correlations between non-polyadenylated and polyadenylated samples for the raw cDNA reads showed large variation across serovars. The extrema of correlation values ranged from R^2^ = 0.478 (LII samples) to R^2^ = 0.704 (LIC samples) (Fig 4a). The correlation values between the non-polyadenylated and polyadenylated samples evaluated with cDNA FL reads showed slightly lower values for all samples, except for LII (R^2^ = 0.524) (Fig S3).

**Fig 3.**
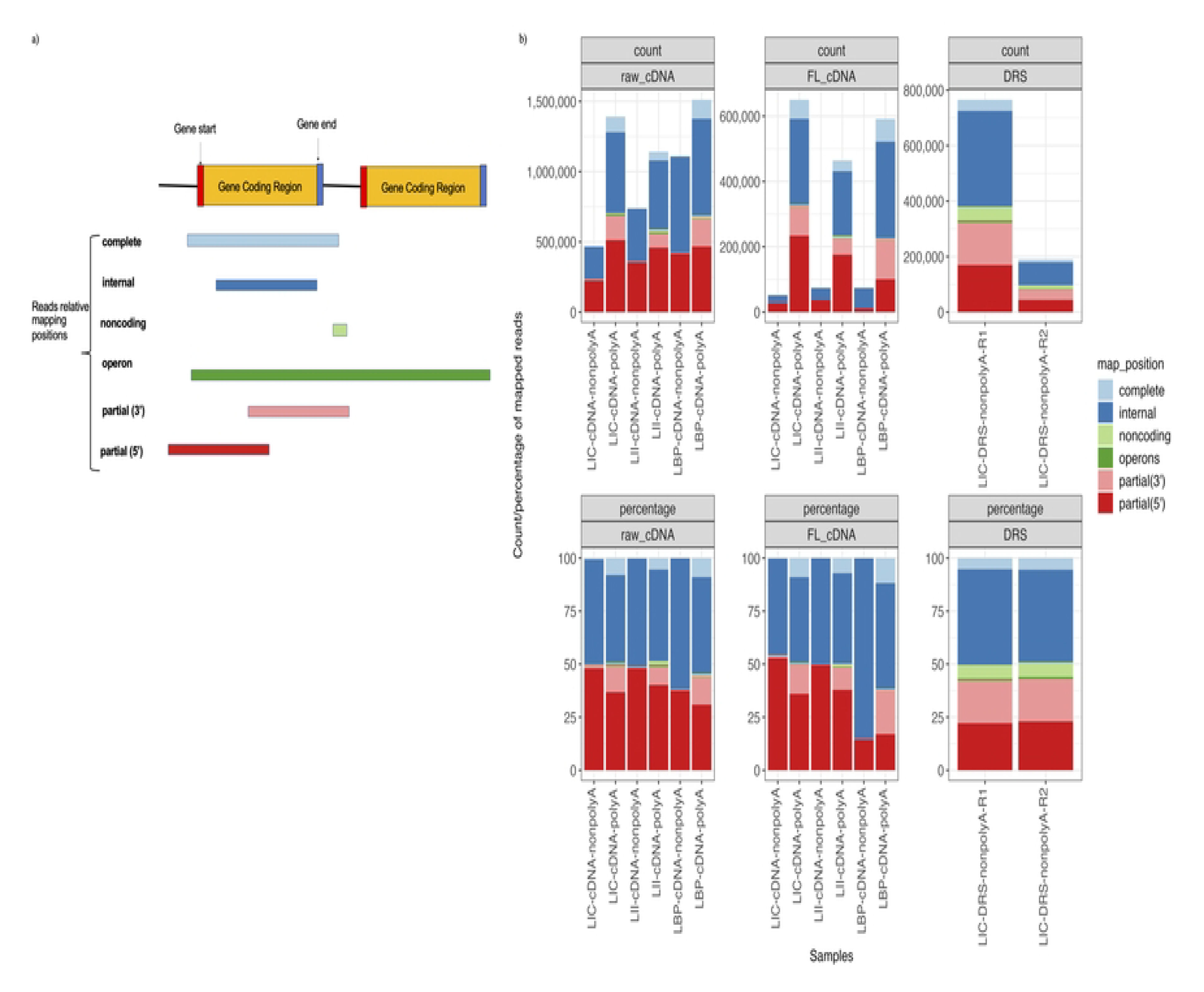
Relative mapping positions and coverages. (a) Diagram showing the six categories of reads mapping positions relative to their closest annotated coding regions in the reference genome. (b) The number and percentage of reads mapped to each mapping position category are listed in panel (a).

**Fig 4.**
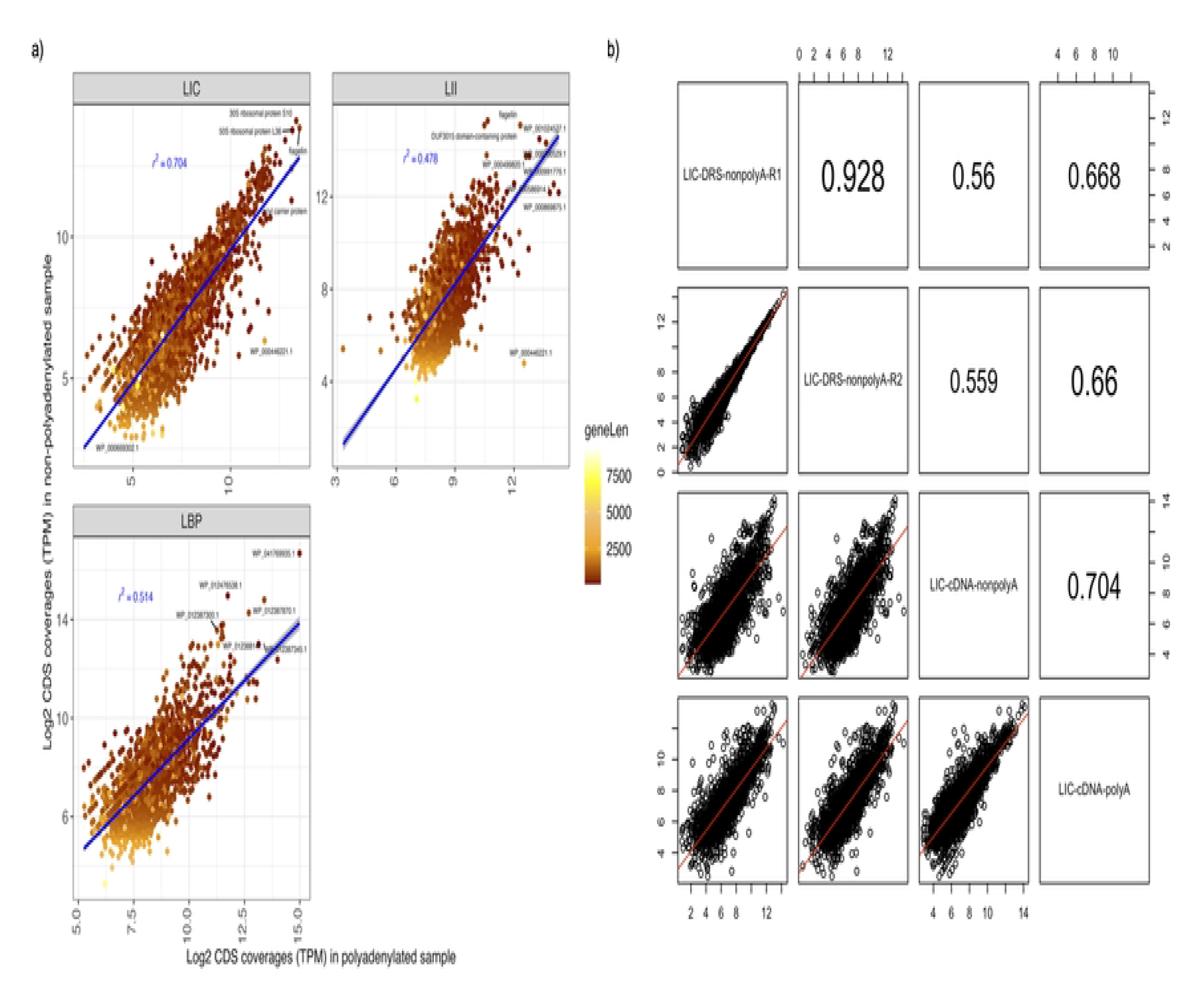
Correlation between coverages of mapped coding regions. Correlations in the mapped annotated coding regions’ coverages (in log2 of transcript per million, TPM) between the polyadenylated and non-polyadenylated raw sequenced cDNA samples and between the LIC samples sequenced with the cDNA and DRS protocols. (a) Correlations in the mapped coding regions coverages between the non-polyadenylated cDNA (y-axis) and the polyadenylated cDNA (x-axis) samples from each serovar. Each dot represents an annotated coding region mapped by both the non-polyadenylated and polyadenylated samples in each serovar to their corresponding reference genome. The color of each dot is scaled by the length (in base pairs) of the corresponding coding regions. (b) Correlations in the annotated coding regions coverages between the cDNA and DRS sequenced LIC samples. The numbers in the upper right diagonal boxes in the matrix are the R^2^ values for the corresponding comparisons between samples labeled in the diagonal boxes directly below or left of the R^2^ value.

On average, over 5% of the DRS reads were categorized as “complete” mapping reads, which was higher than the percentage of LIC’s non-polyadenylated cDNA reads (Fig 3b). The largest “complete” mapping DRS read also covered the annotated coding region of 23 ribosomal RNAs with 2,958 bp long. In addition, DRS reads mapped more reads on average to the “partial (3’)” positions than both raw and FL cDNA reads (20%) (Fig 3b).

We further examined the correlation between the coding regions coverage of the two sequencing protocols, cDNA, and DRS (Fig 4b) of LIC samples. The correlation between the LIC-DRS reads, and each of the polyadenylated and the non-polyadenylated LIC cDNA reads showed a medium to low correlation between the reads from the two sequencing methods, where the correlation between the LIC-DRS reads. The correlation with the polyadenylated LIC cDNA reads was slightly higher (R^2^ ∼ 0.66) than the correlation between LIC-DRS reads and the non-polyadenylated LIC cDNA reads (R^2^ ∼ 0.56).

To evaluate transcripts of the annotated coding regions that underwent potential posttranscriptional polyadenylation, we identified the top 50 most sequenced transcripts across the different serovars (Table S1). Many of these highly sequenced transcripts were identified from both cDNA (non-polyadenylated and polyadenylated) samples, and DRS reads. These transcripts included transcripts encoding for flagellin/flagellar structural proteins, response regulators, ribosomal subunits, RNA transcription factors, catabolism pathways-related proteins, and surface lipoproteins. A total of 25 out of the 50 identified transcripts were present in all the LIC samples sequenced by the two different ONT sequencing protocols.

### Identification of operons

Operons were identified as reads covering more than one annotated coding region and accounted for less than 1% of the reads across all samples. The non-polyadenylated cDNA samples had fewer operon mapping reads, and fewer unique operon transcriptional units (composed of coding regions of different genes) than their polyadenylated counterparts in both raw (Fig S4a) and FL cDNA (Fig S5) reads. On the other hand, the total number of operon mapping reads was larger in the LIC-DRS samples than in the raw LIC cDNA samples (Fig S4a), regardless of polyadenylation. Approximately less than 1% of the operon reads mapped to more than five coding regions, and most operon mapping reads covered exactly two annotated coding regions (Fig S4a). Many of the identified operon transcriptional units aligned to the same and opposite strands of their mapped annotated coding regions (Table S2).

The majority of the distinct operon transcriptional units identified in this study (913) were uniquely identified from samples of a single serovar (Fig S4b). Only 15 of these were identified across all serovars. Some operon transcriptional units (102) were shared only between the samples of pathogenic serovars LIC and LII (Fig S4b). Raw cDNA reads identified the largest number of unique operon transcriptional units, followed by the number of the DRS and FL cDNA ones (Fig S4c). Almost all operon transcriptional units identified in the FL cDNA samples were also identified from the raw cDNA samples. More than half of the operon transcriptional units identified from samples of raw cDNA and the DRS datasets overlapped with each other (Fig S4c).

### Identification of unannotated genomic regions

A total of 1,113 unique unannotated genomic regions were identified across all samples (Table S3). After further sequence querying of these unannotated coding regions in the RNA database Rfam, we found that only a small portion of these sequences was previously described in *Leptospira* or in other bacterial genomes. Some of the sequences of the unannotated coding regions have been previously described in the genomes of other *Leptospira* strains, such as the coding regions for the ncRNA, Lig thermometer (which was identified either between the LigA and LigB or next to LigA coding regions), and sRNA molecules 30_29 (which were identified between the coding regions of leucine-tRNA ligase and ABC transporter ATP-binding protein). Other sequences of the identified unannotated coding regions have not been described previously in a *Leptospira* genome but in the genomes of other bacteria, including those identified as part of the CRISPR direct repeats elements, riboswitches coding regions, and ncRNA Flavo-1 (which were first identified in Flavobacteria and later described in multiple other bacterial genomes) in the LIC and LBP samples. In general, more unannotated coding regions were identified from the genomes of the pathogenic *Leptospira* (Fig S6a). Most of these unannotated genomic regions were observed in the polyadenylated cDNA sequencing and DRS methods, and only a small number of these regions were observed in the non-polyadenylated cDNA samples (Fig S6a). In addition, we found that more than half of these unannotated coding regions identified were shared between the two pathogenic serovars (Fig S6b).

### Evaluation of polyadenylated transcripts

ONT direct cDNA’s library preparation protocol uses poly(A) tail capture from an mRNA molecule using an oligo-dT primer (VNP primer) followed by a second reverse transcription step, which transcribes the first cDNA strand after the strand switching step. Therefore, reads with both poly(A) and poly(T) tails will be present in the sequenced cDNA libraries. We used the Tailfindr software to estimate poly(A) tail length and the distribution of read types (identified as poly(A) tail, poly(T) tail, or invalid) (Fig 5a) for cDNA sequences. Invalid reads, in this case, are reads with no nanopore primer identified with high confidence. Since reads were sequenced from 5’ to 3’ using the cDNA library, the sequencing process frequently terminates before it reaches the end of the poly(A) stretch, thus only reads identified with full-length poly(A) tail were used to estimate the tail length. As expected, the polyadenylated samples had relatively longer median ‘A’ and ‘T’ stretches (∼ 28 bp for A tails and ∼ 57 bp for T tails) compared to those of non-polyadenylated ones (∼ 16 bp for A tails and ∼ 22 bp for T tails) (Fig 5a). In the DRS protocol without polyadenylation, the native RNA molecules with native poly(A) tails are the only poly A stretches with their lengths estimated. We used two popular poly(A) tail estimation software Nanopolish and Tailfindr, to estimate the tail lengths on each read sequenced with the DRS protocol. However, we observed only a moderate correlation (R^2^= 0.59) between the tail lengths estimated by the two software. Nanopolish reported a median tail length ∼9 bp and Tailfindr reported a median tail length of ∼15 bp (Fig 5b).

**Fig 5.**
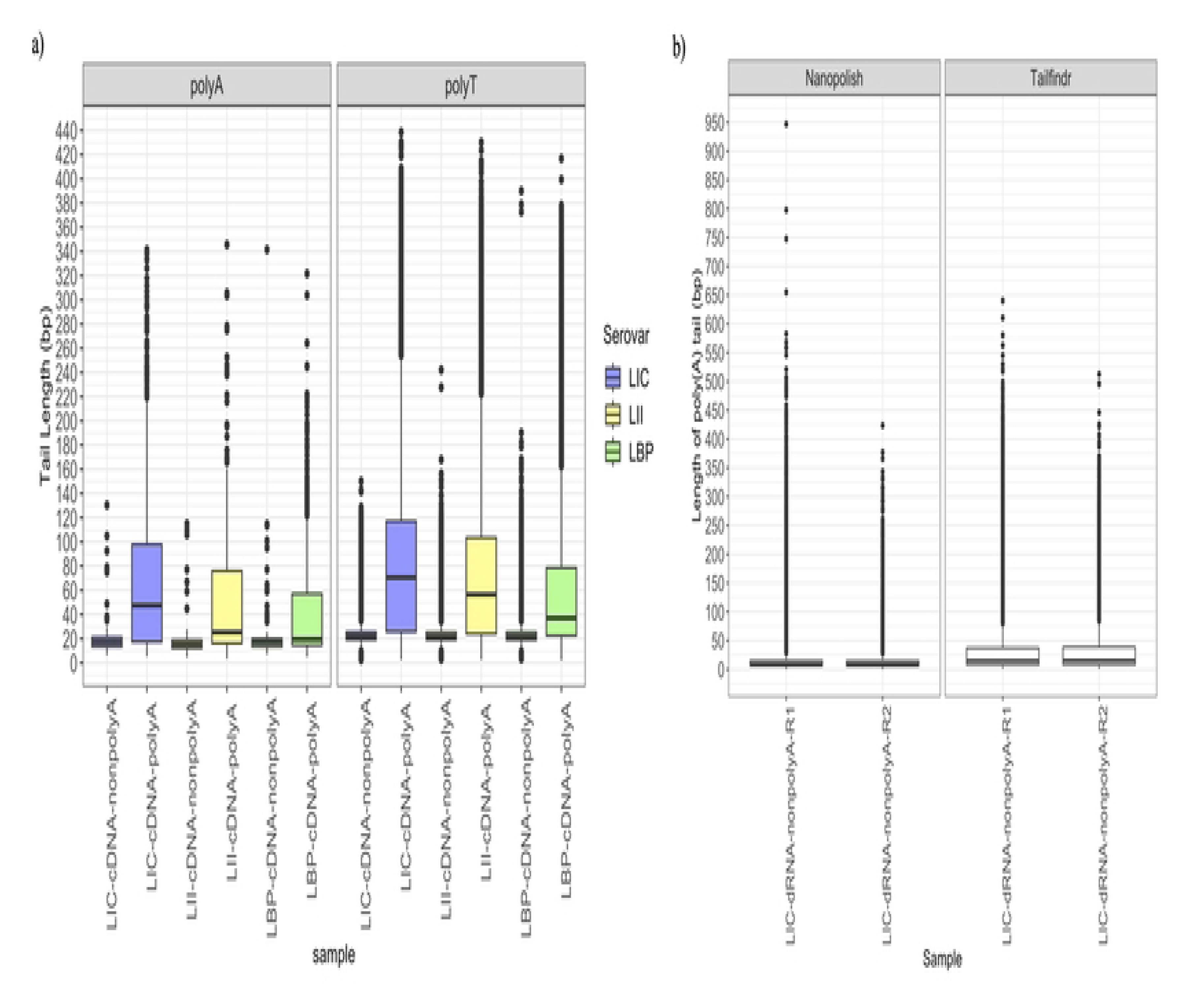
PolyA tail assessment for direct cDNA and DRS reads. The estimated tail length (poly(A) and poly(T)) for the direct cDNA samples is presented in (a). Two different software were used to evaluate poly(A) tail length in DRS samples. The estimated tail lengths reported by the two software are shown in (b).

As depicted in Fig 6a, ONT sequencing methods have the potential to prime and sequence the fragmented transcripts with truncations at the homopolymers of adenine (A) regions within the transcribed coding sequences, causing a potential drop in coding region coverages after the presence of the homopolymers of adenine (A) regions at the 3’ end. This process might especially be prominent in organisms with AT-rich genomes, such as *Leptospira.* To assess the bias introduced due to the presence of homopolymers of A within the transcripts, we analyzed the average relative coverage (see Materials and Methods) of bases upstream and downstream of all identified homopolymers of A regions (position 0) from the annotated coding regions (Fig 6b). The averages of the relative coverages of each position flanking the identified homopolymer of A regions (across all identified homopolymer of A regions from all samples) were calculated and reported in Fig 6b. With the presence of enzymatic poly(A) tail attached to the 3’ end of all sequenced transcripts, polyadenylated cDNA samples were found with no decrease in the relative coverage after the presence of a homopolymer of As. On the other hand, by capturing only the native poly(A) attached to the sequenced transcripts, the non-polyadenylated raw cDNA samples show a slight drop in average 3’ relative coverage after the presence of a homopolymer of A. This implies biases in capturing fragmented transcripts truncated at homopolymers of A regions during library preparation (Fig 6b). However, the average 3’ end coverage drop after the homopolymer of A regions was steeper within the FL samples, especially in the non-polyadenylated ones. In addition, the average relative coverages flanking the homopolymers of A regions in the LIC-DRS samples were also largely dropped at the 3’ end after the presence of the homopolymer of A regions (Fig 6b).

**Fig 6.**
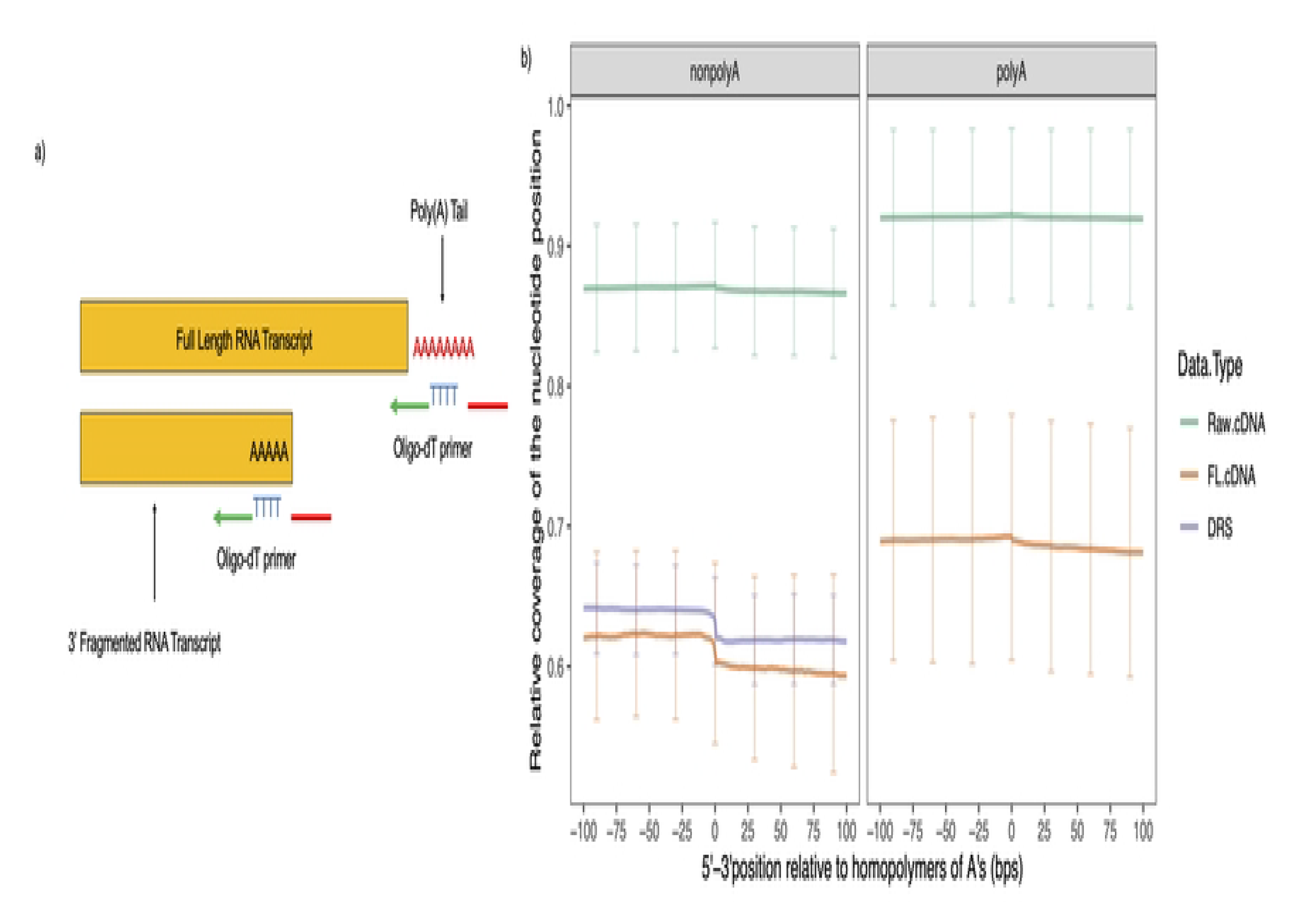
Enrichment in sequencing transcripts with 3’ truncation at a homopolymer of A regions. (a) Oligo-dT primers in both cDNA and DRS protocols may pick up RNA transcripts fragmented at homopolymer of A regions within the coding sequences to initiate transcription during library preparation. (b) Average relative coverages for nucleotides flanking to the homopolymer of A regions (>= 5bps) in the raw/FL cDNA and DRS samples. Relative coverage at each nucleotide position was calculated in relation to the maximum coverage within the annotated coding region where the homopolymer was identified.

## DISCUSSION

In this study, we utilized ONT’s long read sequencing technology to evaluate *Leptospira* transcriptome, identify potential coding regions of multiple RNA molecules that have not been previously described, and determine operon transcriptional units encoding for transcripts of multiple genes. Our analysis suggested that some of the sequences containing homopolymers of adenine bases at the 3’ end of RNA molecules may in fact, be the result of posttranscriptional polyadenylation described in prokaryotes [34].

For the unannotated coding regions identified in this study, most of them had never been reported previously in *Leptospira* transcriptome. Many of the relative positions of these regions were consistent in the identities of their neighboring coding regions across the reference genomes of different pathogenic serovars analyzed in this study. This consistency could be the result of large horizontal gene transfer with the involvement of multiple genes during *Leptospira’s* adaptation mechanism [5]. Furthermore, the majority of the unannotated coding regions were not detected in the nonpathogenic *Leptospira*. In addition, the relative positions of the unannotated coding regions identified in the nonpathogenic *Leptospira* were not consistent with those identified in the pathogenic strains, suggesting potential virulence related functions for the RNA molecules transcribed from these unannotated regions. More interestingly, many operon transcriptional units identified from the pathogenic strains were also not observed in the nonpathogenic strain, suggesting different gene regulatory mechanisms present in the pathogenic and nonpathogenic *Leptospira* species. Carefully designed experiments are needed to unravel the role of these regions in *Leptospira* virulence and pathogenesis. Although we observed some evidence for variations in RNA expression between strains analyzed in this study, the data should be interpreted with caution considering the inherent issues associated with the ONT sequencing methods [32,35–37]. We propose further confirmation using hybrid sequencing methods and carefully designed experiments with appropriate replicates.

The advantage of using ONT’s direct long read sequencing protocols is its ability to sequence the full-length RNA transcript per read with no theoretical read-length limitations or amplification biases [38], thus allowing the detection of different isoforms and intermediate structures of RNA transcripts [36,39]. In addition, with direct sequencing of RNA molecules and reverse transcribed cDNA molecules, ONT’s long read sequencing technology can assess the presence of posttranscriptional modification structures in the sequenced libraries, including the presence of poly(A) tail [32,40]. In our initial experiments with ONT’s RNA-Seq technologies, despite skipping the polyadenylation step required for sequencing of prokaryotic mRNA, the observation of obtaining a moderate yield of transcript reads was both interesting and intriguing. Subsequent comparisons were made using pathogenic and nonpathogenic *Leptospira* using RNA transcripts sequenced with and without polyadenylation. While there were no differences in quality and read length of non-polyadenylated vs. polyadenylated reads, the sequencing yield in the non-polyadenylated samples and the percentage of reads identified as FL reads were significantly smaller in the non-polyadenylated samples. This is expected because only transcripts with poly(A) tail attached were expected to get sequenced in the non-polyadenylated library. In DRS, a higher percentage of mRNA molecules was sequenced, and a similar number of transcripts (from distinct annotated coding regions) was sequenced in comparison to the polyadenylated direct cDNA samples, but the average read length was shorter. This could be due to the DRS’s capability of capturing more intermediate RNA transcripts in the process of degradation [41]. However, our attempt to sequence RNA by DRS after the polyadenylation step generated only less than ∼0.025 Gb of data and these were not enough for a comparative analysis (data not shown in this study). We believe that this could be a result of pore blocking by the RNA secondary structures during the sequencing process [35,37]. Surprisingly, DRS samples sequenced without polyadenylation generated a significantly higher sequencing yield (with almost 40 times more data on average) than the DRS samples sequenced with polyadenylation. The non-polyadenylated DRS sequenced samples had a lower percentage of their sequenced reads mapped to the reference genome than the cDNA sequenced ones. This suggests that the reads suffered the effects of higher error rate during sequencing for the DRS protocols [32], which was further confirmed by the lower mapping identity evaluated from the DRS samples’ alignments.

Although largely recognized and described in eukaryotes, the presence and function of polyadenylation resulting in 3’poly(A) tails in prokaryotic transcriptome has not been well characterized [8,10,22,33,34]. Previous studies of prokaryotic polyadenylation have found that, unlike the eukaryotic poly(A) tails, which are mainly attached to the 3’end of mRNA transcripts [42,43], the polyadenylation sites in the prokaryotic transcriptome can attach to mRNA, rRNA, tRNA, tmRNA, and sRNAs molecules [44]. Due to their role in RNA degradation mechanisms, prokaryotic poly(A) tails can be identified not only at the end of a full-length primary transcripts, but also in the cleaved intermediate transcripts processed by endonucleases [45,46]. The presence of a poly A polymerase was described in *E. coli* as early as 1962 and the gene coding regions for poly(A) polymerase (PAPI), *pcnB,* was later identified in *E. coli* as the major contributor for polyadenylation of RNA transcripts [47,48]. Later, this gene was identified in a limited number of gram-negative bacterial genomes [49,50]. Interestingly, the differential expression activity of the *pcnB* gene has been identified in a previous transcriptome study on pathogenic *L. interrogans* when cultured under different conditions [8]. We also identified that many of the highly sequenced transcripts in the non-polyadenylated samples are consistent across serovars and species analyzed, suggesting the presence of shared mRNA profiles potentially regulated by inherent polyadenylation across *Leptospira* species. It is noteworthy that many of these highly sequenced transcripts in the non-polyadenylated samples were also found regulated by the polyadenylation in *E. coli*. These include transcripts encoding for membrane proteins, motility proteins, and RNA metabolism proteins [34,51].

In theory, ONT’s direct RNA-Seq protocols could sequence a complete transcript in different isoforms [32,36], and the relative mapping positions of each read to the coding regions should reflect fragmentated gene transcripts. We found that compared to the polyadenylated samples, only a small percentage of reads sequenced without polyadenylation (in both cDNA and DRS samples) was able to align with the complete coding regions of their mapped reference genome. Even for the polyadenylated samples, the majority of the sequenced reads were identified mapping only partially with the complete annotated coding regions. This suggests that a large percentage of RNA molecules in different *Leptospira* transcriptomes are potential intermediate transcript fragments in the process of degradation. In addition to the reads fully covering the annotated coding regions, samples sequenced without polyadenylation were also reported having less percentages of reads mapped to the 3’ end of the mapped genes’ coding regions compared to the polyadenylated sequenced samples. This could be the result of the non-polyadenylated samples only capturing transcripts degraded from the 3’ end with the assistance of poly(A) tail. However, the observation with the lack of 3’ partially mapped reads in the non-polyadenylated samples were only observed within the direct cDNA sequenced samples. This could be due to DRS’s ability in capturing a complete RNA molecule without polyadenylation [41]. Our data suggested the presence of gene transcripts potentially underwent degradation with the assistance of poly(A) tails.

Although our study provided the opportunity to explore *Leptospira*’s transcriptomic landscape and RNA structures without enzymatically polyadenylating samples, it also presents a few limitations. Firstly, in addition to sequence transcripts possessing a true enzymatically added poly(A) tail, transcripts truncated at the homopolymers of A regions during library preparation are also captured, which can be a significant proportion of RNA profiles of AT rich *Leptospira* genome. While this method is advantageous to study post-transcriptional polyadenylation, sequencing of transcripts truncated at the homopolymer of A regions could be the source of bias for making conclusions in RNA composition and differential gene expression studies. Secondly, we observed that algorithms (Pychopper) used to identify FL cDNA reads could falsely identify sequences in the middle of a raw cDNA read with similar nucleotide identity as the reverse primers. This misidentification of reverse primer will falsely classify a non-FL read as a FL cDNA read and trim a raw cDNA read in the middle of the sequenced transcripts from the misidentified primer. Thus, DRS and the FL cDNA reads identification process should be taken when sequencing prokaryotic transcriptomes without polyadenylation. These biases could not only affect the conclusion about the poly(A) tails in the prokaryotic transcriptome but also will bias the relative profiles of gene expressions. We also observed that poly(A) tail length estimation software reported diverged estimation for the poly(A) tail length of the non-polyadenylated DRS samples; thus, we should also be cautious about the tail length estimation from the non-polyadenylated DRS samples. However, the range of poly(A) tail length estimated from the non-polyadenylated cDNA samples (∼16 bps) and one of the tail length estimation software for the DRS samples (∼15 bps using Tailfindr) is consistent with the previous estimation for prokaryotic poly(A) tail *in E.* coli (14-60 bps) [34]. Since only one software is currently developed for direct cDNA tail estimation, we could not assess the accuracy of poly(A) tail estimation in the direct cDNA samples in this study.

In conclusion, by using the ONT’s direct cDNA and RNA sequencing protocols in this study, we provide a preliminary comparison and an opportunity to evaluate the *Leptospira* transcriptome. ONT’s ability to detect and sequence prokaryotic RNA molecules without polyadenylation opens an opportunity to explore potential post transcriptional polyadenylation and its effects in prokaryotic gene regulation. Further evaluation of the unannotated genomic regions and operon transcriptional units discovered in this study will unravel the complex virulence, pathogenesis, and host adaptation mechanisms of *Leptospira*, an understudied spirochete of public health significance.

## MATERIALS AND METHODS

### Bacteria

We used two pathogenic *L. interrogans* serovars (Copenhageni [LIC] and Icterohemmorhagiae [LII]) all belonging to the same serogroup (Icterohemmorhagiae) and one nonpathogenic serovar of *L. biflexa* (serovar Patoc [LBP]). Bacteria were grown in Ellinghausen-McCullough-Johnson-Harris (EMJH) liquid medium, supplemented with Difco *Leptospira* Enrichment EMJH (Becton Dickinson, Sparks, MD, USA) at 29 °C and a 4-day old culture was used for RNA extractions. RNA from each strain was extracted using a commercial kit (miRNeasy Mini Kit, Qiagen, Hilden, Germany) following the manufacturer’s protocol. After quantity and quality assessment using Qubit (Thermo Fisher, Waltham, MA, USA) and NanoDrop (Thermo Fisher, Waltham, MA, USA), the RNAs were further cleaned and concentrated using RNA Clean and Concentrator (Zymo Research, Irvine, CA, USA). The final assessment of RNA quality and quantity was done using Bioanalyzer (Agilent, Santa Clara, CA, USA).

### RNA sequencing using the direct cDNA sequencing method

We used Direct cDNA Sequencing Kit (SQK-DCS109 Oxford Nanopore Technologies, Oxford, United Kingdom) for the library preparation following the manufacturer’s instructions. One aliquot of RNA from each *Leptospira* serovar was processed with the addition of poly(A) tail using *Escherichia coli* poly(A) Polymerase Kit (New England Biolabs, Massachusetts, USA) following the protocols from the manufacturer. Before loading the samples in the flow cells, the polyadenylated and non-polyadenylated samples underwent strand switching process, barcoding for multiplex sequencing, and adapter ligation. Up to six samples were multiplexed in a single run. The sequencing process was conducted in MinION Sequencing platform (Oxford Nanopore Technologies, Oxford, United Kingdom) for 48 hours with bias voltage set to −180 mV.

### RNA sequencing using the direct RNA sequencing method

We sequenced RNA from two replicates of LIC *serovar* without the polyadenylation step using Direct RNA Sequencing Kit (SQK-RNA002, Oxford Nanopore Technologies, Oxford, United Kingdom). The library preparation was done following the manufacturer’s instructions. The library preparation process included attaching the RNA to the RT adapter and the sequencing adapters to the ends of the RNA. Since the Direct RNA Sequencing Kit does not allow multiplexing, each sequencing was carried out in a single MinION flow cell for 24 hours with bias voltage set to −180 mV.

### Raw sequence quality check, basecalling, and demultiplexing

The quality of the raw cDNA and DRS Fast5 files were first assessed using MinIONQC v.1.4.1 [52]. After quality check, the sequences were base called using Guppy v. 4.4.2 (https://community.nanoporetech.com) with the option “—config dna_r9.4.1_450bps_hac.cfg” for the cDNA samples and option “—config rna_r9.4.1_70bps_hac.cfg” for the DRS samples in to the FASTQ format. FASTQ files of cDNA reads were further demultiplexed using Guppy’s “guppy_barcoder” function with the option “—barcode_kits EXP-NBD104”. After demultiplexing, the barcoding summary produced with “guppy_barcoder” was used to separate for each sample multiplexed cDNA Fast5 files into the demultiplexed Fast5 files using the “demux_fast5” function provided by ont_fast5_api (https://github.com/nanoporetech/ont_fast5_api). The separated Fast5 files for each individual cDNA sample were then re-basecalled with the same configuration as above. MinIONQC was used again to assess the quality of each individual cDNA sample.

### FL reads identification and processing

FL reads from the cDNA raw sequence reads were identified using Pychopper (v. 2.7.1) (https://github.com/epi2me-labs/pychopper) following the suggested FL identification protocol designed for bacterial RNA-Seq from a previous study [37]. The raw reads were first processed with the default Pychopper parameters to autotune the cutoff values from the input cDNA reads of each sample. The reads that could not be identified as FL reads in the first round (unclassified reads) went through a rescue process using the parameter (-x rescue) in the second round of processing with a direct cDNA specific rescue option (DCS109) turned on. The processed reads and the rescued reads from both rounds of processing were concatenated together to create the FL reads dataset for each cDNA sample.

The average hit qualities of the SSP and VNP primers from each raw cDNA reads were obtained from the Pychoppers’ summary reports (obtained with the “-A” option specified) and calculated using a custom R script [53]. The lengths of the subsequences trimmed off from each end of the raw cDNA reads to the position of the identified SSP and VNP primers were obtained from comparing the original read length to the read trimmed positions reported in the Pychoppers’ summary report. Since the cDNA library preparation protocol generates two strands of cDNA molecules starting from the 3’ end poly(A) stretches of the targeted native RNA molecule, the first cDNA strand is the reverse complemented read of the native RNA molecule with the VNP primer at the beginning of the read and the reverse complemented SSP primer at the end (VNP, -SSP). The second cDNA strand is generated after strand-switching possessing the SSP primer at the beginning of the sequenced read and the reverse complemented VNP primer at the end (SSP, -VNP). Therefore, for the SSP primers (SSP) or the VNP primers (VNP) identified at the beginning of each read: trimmed-off length = trimmed position -a, where ‘a’ indicates the start position of the read (0), and for the reverse-complemented SSP primers (-SSP) or the reverse-complemented VNP primer (-VNP) at end of each read: trimmed-off length = length of the raw cDNA read – trimmed position + 1. The average trimmed off lengths obtained from either SSP or VNP primers of all identified FL cDNA reads of each sample were calculated using a custom R script.

### Mapping to the reference genome

Raw and FL cDNA reads were mapped to their corresponding reference genomes using Minimap2 v. 2.17 [54], separately, with the options “-ax splice -p 0.99 –MD –cs -Y”. DRS reads were mapped with the options “-ax splice -uf -k14 -p 0.99 –MD –cs -Y”. Serovar specific reference genomes were used for mapping, where *L. interrogans* serovar Copenhageni str.

Fiocruz L1-130 (GCF_000007685.1) was used for LIC samples, *L. interrogans* serovar Icterohaemorrhagiae str. Langkawi (GCF_014858895.1) was used for LII samples and *L. biflexa* serovar Patoc str. ‘Patoc 1 (Paris)’ (GCF_000017685.1) was used for the LBP samples. Each sample’s alignment files in binary format (BAM) were first sorted and indexed using SAMTools v.1.10 [55], and converted into BED format for downstream analysis using the “bamtobed” function in BEDTools v. 2.30.0 [56] with the “-cigar -tag NM” options. We further obtained the mapping identities of all reads in each sample using a custom python script with the formula (1-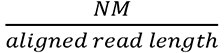* 100) [37], where NM is the number of mismatches and gaps reported by Minimap2 and the aligned read length was reported as the sum of M(atch) and I(nsertion) characters obtained from each read alignment’s CIGAR (Compact Idiosyncratic Gapped Alignment Report) string.

To determine the closest coding region of each mapped read and the relative mapping positions of the read to the coding region, we used a custom python script to compare the mapping positions (mapping start & mapping end) of each read to the reference genome to the annotated coding regions provided by the corresponding reference genome’s GFF file. We further labelled read alignments by the 2D structure reads [37] by identifying reads mapped with the presence of N (intron) in each read alignments’ CIGAR string.

### Evaluation of homopolymer of A regions

The method to assess homopolymer of A region’s coverage was adapted from a previous study [57]. Genomic regions with at least 5 consecutive A’s in the annotated coding regions were identified from all the reference genomes used in this study with a custom python script. The coverage of individual bases that were located 100bp up-and down-stream of the identified homopolymer of A regions were obtained using the “coverage” function in BEDTools v. 2.30.0 [56] with the option “-d”. For FL and DRS reads, the coverages of each base flanking to the homopolymer of A regions were determined in a strand-specific manner with the option “-s”. The relative coverage of each base flanking to the homopolymer of A regions were calculated by using the absolute coverage of each base divided by the maximum coverage of the coding region where the homopolymer was identified from. The maximum coverage of each coding region and the relative coverages of each base were determined by a custom python script.

### Poly(A) tail length estimation

The length of the poly(A) tail for the cDNA sequenced samples was calculated using Tailfindr v 1.0 and the length of the poly(A) tail for the DRS sequenced sampled was calculated using both Tailfindr v 1.0 [58] and Nanopolish v 0.13.2 [59], where the latter is a software designed specifically for DRS sequenced samples.

### Unannotated genomic regions identification

All reads that were identified mapping to the “noncoding” positions in the previous analysis were extracted and grouped together based on the closest annotated coding regions they were mapped to. Those groups with only one read were filtered out from the downstream analysis. After filtering, the mapped positions with the highest coverage at the 3’ and 5’ ends within each group of reads were obtained as the potential start and end positions of the unannotated coding regions, respectively. Nucleotide sequences of these extracted regions were extracted using the “getfasta” function in BEDTools v. 2.30.0 [56]. The regions with duplicated sequences and sequences larger than 200,000 bp were filtered. Remaining sequences were queried in the Rfam database [60] for further annotation.

### Operon transcriptional units identification

Reads that were identified to cover more than one annotated coding regions were identified as operon reads. To avoid the reads partially covering the coding regions are only the transcribed 5’ or 3’ UTR sequences overlapped with the flanking coding regions, only the annotated coding regions covered at least 90% by the mapped reads were included. In the end, the operon reads covering the same annotated coding regions were identified as the unique operon transcriptional units.

### Correlation of coding region coverages

The number of reads mapped to each annotated coding region was obtained using HTSeq v. 0.9.1 [61] with the option “—nonunique all –type=CDS”. This option counts for all reads, even the ones that were mapped to more than one coding region, contributing to the coverage of the mapped coding region the reads belong to. The number of reads mapped to each coding region was then normalized into TPM (transcripts per million) using a custom R code. Correlation analyses between the normalized coding regions coverages were performed and determined using the “lm” function in the R “stats” package [53].

## Acknowledgement

We thank The Interdisciplinary Disease Ecology Across Scales program (Funded by the National Science Foundation under Grant No. DGE-1545433) at The University of Georgia for providing funding and training to R.X when completing this manuscript, The University of Tennessee, College of Veterinary Medicine for Start-up funds for S.R to set up *Leptospira* research laboratory, and The University of Georgia, Office of Research for Start-up funds for L.C.M.S.

## Supporting Information

Supplementary_figures.docx

TableS1.top50_most_highly_mapped_CDS.xlsx

TableS2.operon_structures.xlsx

TableS3.unannotated_coding_regions.xlsx

## Data Availability

All direct cDNA sequenced samples are available under bioproject: PRJNA949181, with biosample accession: SAMN33927558, SAMN33927559, SAMN33927560, SAMN33927561, SAMN33927564, SAMN33927565. All direct RNA sequenced samples are available under bioproject: PRJNA949189, with biosample accession: SAMN33933649 and SAMN33933650. All scripts for mapping analysis are available in the github repository: https://github.com/rx32940/Lepto_transcriptome_ONT_cDNA.

